# Utility of cellular imaging modality in subcellular spatial transcriptomic profiling of tumor tissues

**DOI:** 10.1101/2024.07.25.605221

**Authors:** Xiaofei Song, Xiaoqing Yu, Carlos M Moran-Segura, G Daniel Grass, Roger Li, Xuefeng Wang

**Author notes:** To whom correspondence should be addressed: Xuefeng Wang, Moffitt Cancer Center,; Roger Li, Moffitt Cancer Center,.

## Abstract

Spatial transcriptomics (ST) technologies, like GeoMx Digital Spatial Profiler, are increasingly utilized to reveal the role of diverse tumor microenvironment components, particularly in relation to cancer progression, treatment response, and therapeutic resistance. However, in many ST studies, the spatial information obtained from immunofluorescence imaging is primarily used for identifying regions of interest, rather than as an integral part of downstream transcriptomic data interpretation. We developed ROICellTrack, a deep learning-based framework, to better integrate cellular imaging with spatial transcriptomic profiling. By examining 56 ROIs from urothelial carcinoma of the bladder (UCB) and upper tract urothelial carcinoma (UTUC), ROICellTrack accurately identified cancer-immune mixtures and associated cellular morphological features. This approach also revealed different sets of spatial clustering patterns and receptor-ligand interactions. Our findings underscore the importance of combining imaging and transcriptomics for comprehensive spatial omics analysis, offering potential new insights into within-sample heterogeneity and implications for targeted therapies and personalized medicine.

## INTRODUCTION

Spatial transcriptomics (ST) technologies, such as GeoMx Digital Spatial Profiler (DSP) by Nanostring and Visium platform by 10x Genomics, has seen widespread use in the cancer genomics community for their unparalleled ability to elucidate spatial heterogeneity and the complex interactions within the tumor-immune microenvironment. Multiplex immunofluorescence (mIF) staining, the technique enabling the simultaneous detection of multiple biomarkers on the same tissue slides, is a critical preparatory step before employing ST analysis. This approach provides a comprehensive overview of cellular composition within the tissue, serving as a crucial check for cellular heterogeneity, the spatial distribution of different cell types, and the presence of specific cellular markers. The information collected from H&E and mIF staining is then utilized to accurately identify regions of interest (ROIs) for targeted transcriptome assays based on NGS read-outs. For example, the commonly used Morphology Marker Kit for studying solid tumors in the DSP contains fluorescent biomarkers: PanCK (for tumor cells), CD45 (for immune cells), SYTO13 (for visualizing nuclei), and customized biomarkers (e.g. CD20 for identifying B cells). Of note, DAPI, a commonly used fluorescence dye for nuclei staining, is not used in the DSP system. We have observed that in most applications of GeoMX DSP, spatial information derived from multiplex immunofluorescence imaging has been predominantly utilized for identifying targeted regions for molecular profiling without fully integrating this rich spatial context into downstream transcriptomic data analysis.

Although ROI-based gene expression data offer detailed insights into a specific region at the subcellular level, the bioinformatics pipelines that follow often treat these spatially-resolved datasets similarly to how one might approach RNA expression data analysis with bulk tissues.

Recognizing this missed opportunity at the subcellular level, we developed a comprehensive analytical framework that employs deep learning to fully leverage the intricate spatial information provided by in situ imaging. This approach enhances the utility of cellular imaging modalities in spatial transcriptomic profiling, allowing for a more detailed understanding of the spatial distribution of gene expression within tumor tissues. Our study focuses on urothelial carcinoma of the bladder (UCB) and upper tract urothelial carcinoma (UTUC), analyzing 56 ROIs with the aim of bridging the gap between spatial transcriptomic data and cellular imaging, thereby offering a novel perspective on tumor spatial analysis.

By integrating advanced image-based analysis with ROICellTrack, our framework demonstrates the potential to provide novel insights into cancer cell populations, cell-level morphological features, and the complex interplay between tumor cells and their microenvironment (Figure A). This integration not only enhances our understanding of tumor heterogeneity but also opens up new avenues for targeted therapies and personalized medicine. Through this innovative approach, we underscore the importance of combining spatial transcriptomics with cellular imaging to achieve a comprehensive understanding of tumor biology at the subcellular level.

## RESULTS AND DISCUSSION

To demonstrate the capabilities and advantages of integrating spatial information into gene expression analysis, we analyzed data generated from bladder cancer patients. This investigation involved spatial RNA profiling of tumor specimens from individuals diagnosed with both Upper Tract Urothelial Carcinoma (UTUC) and Urothelial Bladder Cancer (UBC), utilizing the GeoMx DSP platform (Figure B). In this study, we identified six Formalin-Fixed, Paraffin-Embedded (FFPE) samples from concurrent radical cystectomy and nephroureterectomy performed on three patients. We selected and analyzed a total of 56 evaluable Regions of Interest (ROIs) to explore spatial heterogeneity and its association with gene expression patterns in these areas. Each ROI was carefully chosen by the study pathologist to ensure representative coverage of both lower-tract (LW-Tu) and upper-tract (UP-Tu) locations. This selection process also aimed to include a diverse spectrum of the tumor microenvironment: including tumor cores (Tu), stroma (St), tumor-stroma margins (TuSt), and regions with high immune cell infiltrates. These ROIs provide useful data essential for benchmarking the estimation of tumor continent and cellular composition. Multiplex IF was performed on each FFPE specimen with stains for DNA SYTO13, PanCK, CD20, and CD45. The GeoMx DSP platform was used to capture the RNA reads from the ROIs on each FFPE slide.

To facilitate the analysis of image data from DSP ROIs, we created a Python-based toolkit named ROICellTrack. This toolkit streamlines the entire workflow of subcellular image analysis, integrating processes from image preprocessing to cell segmentation and downstream quantitative analysis. As shown in Figure A, the initial step in the ROICellTrack workflow involves the cropping of images to focus on the ROI. While the DSP analytic platform facilitates the batch download of zoomed-in images centered around the ROIs, ROICellTrack enhances this process by further cropping these images to retain only the regions within the ROI that match a predefined shape, typically a circular shape as shown in our study. The main cell identification pipeline was built based on Cellpose, which is a state-of-the-art cellular segmentation model based on U-Net neural network architecture. Cellpose also includes an ensemble of diverse pre-trained models which offers the capability for easy re-training, allowing users to fine-tune models based on their specific datasets. For our analysis, we used the pre-trained model based on TissueNet, which is a large fluorescent microscopy dataset collected from different platforms. Following segmentation, the cells identified within the Region of Interest (ROI) can be visualized using post-segmentation plots. These plots can be presented through a cell mask view, which aids in analyzing cell size by providing a clear representation of each cell’s shape and area. Alternatively, visualization can be achieved through a cell boundary view, which focuses on the outlines of cells, thereby facilitating the examination of cellular compartments and their spatial relationships.

Following cell segmentation, our next steps involve quantifying cell numbers and analyzing the average color intensities within each cell. As a demonstration of this process, we focused on three color channels: green representing PanCK (indicative of cancer cells), red for CD45 (marking immune cells), and blue for DNA. The distribution plot of these three color channels proved highly informative. Notably, the green channel exhibited a bimodal distribution, which serves as a basis for major cell typing, distinguishing between cancerous and non-cancerous cells. The red color channel, representing immune cells, does not display a bimodal distribution as visually inspected, primarily due to the inherent blending of red with other colors within the image and impacted by the overall warm color (like reds and yellows) bias of the input image. To further refine our cell typing, we applied clustering methods, specifically K-means and Gaussian Mixture Model (GMM) clustering, focusing on the green and red channels. Our findings indicated that GMM clustering yielded more definitive results compared to K-means. Because the green channel’s intensity is effectively clustered into groups, it allows us to build a clustering-free method and establish a simple cutoff at an intensity value of 10 to separate the two cell groups. As shown in the right panel of Figure A, utilizing our cell typing strategy, we were able to estimate that out of a total cell count of 535 within the example regions, 258 were identified as cancer cells. Consequently, the proportion of cancer cells in relation to the total cell population is calculated at 48.22%. This figure offers an absolute measure of the cancer cell burden within the sample, presenting a more accurate assessment compared to relative methods dependent only on total color intensities. ROICellTrack is designed to automatically generate detailed morphological features at the cell level. These features encompass R/G/B intensities, the total number of pixels within each identified cell, as well as the cell’s perimeter, area, and circularity. This comprehensive dataset offers a wealth of information for detailed cell-level downstream analysis, allowing researchers to study not just the color properties but also the structural attributes of cells.

To demonstrate the accuracy of ROICellTrack in cancer cell assessment, we compared its performance against a deconvolution-based approach that infers cellular composition based on gene expression levels of cell markers. SpatialDecon is a spatially resolved cell deconvolution algorithm specifically designed for the gene expression modality of GeoMX DSP datasets. Using SpatialDecon, we deconvolved each region of interest (ROI) based on expression signatures representing tumor cells, as well as common stromal and immune cell types within the tumor microenvironment (TME). Our analysis revealed a significant positive correlation between ROICellTrack and the deconvolution method (R = 0.9, P < 2.2e-16), although discrepancies were noted. ROICellTrack accurately identified high tumor purity in Tu cores, moderate levels of tumor cells in TuSt ROIs, and low tumor abundance in St cores. In contrast, deconvolution tended to underestimate tumor purity in tumor regions by misclassifying some cancer cells as immune cells, while overestimating tumor abundance in stromal areas, with discrepancies reaching up to 0.41 (Figure C). Our findings underscore the high accuracy of ROICellTrack, particularly in ROIs with either markedly high or low fractions of cancer cells.

Beyond assessing tumor purity, we further leverage the outputs from ROICellTrack to reveal spatial clustering patterns that cannot be inferred from the sequencing modality alone. As illustrated in Figure B, tumor cells may either distinctly separate from stromal cells or exhibit a blended mixture with them. We hypothesized that these two groups of tumor-stroma margins could foster different levels of cell-cell communication, thereby presenting distinct transcriptomic features. To test this hypothesis, we first systematically evaluated the degree of tumor-stroma mixture by conducting a spatial clustering test (Figure D). The cell distributions on tissue slides were extracted as point patterns, with each cell represented as a dot. The clustering degree was calculated using the cross-K function to compare observed patterns with a reference Poisson Process. The area under the curve (AUC) was used as a continuous score that represents the degree of mixture. A higher score indicates a more mixed pattern. We then ranked the TuSt ROIs by AUC score and categorized them into Separative (n=19) and Mixture (n=7) groups. Dimension reduction analysis with UMAP revealed overall transcriptomic differences among ROIs from different regions. Separative cores were found to be closer to Tu cores on the UMAP plot, whereas Mixture cores formed a separate cluster positioned between tumor and stroma (Figure E left panel). The immune infiltration status was manually examined using H&E and mIF staining by a pathologist. Immune-enriched cores were observed to be positioned separately from immune-depleted cores (Figure E right panel). More interestingly, Mixture cores are frequently recognized as immune-enriched, whereas most Separative cores were identified as immune-low/depleted. Motivated by these observations, we performed a differential gene expression analysis comparing Mixture and Separative TuSt ROIs. We found that Mixture cores exhibit upregulation of EPSTI1, a gene induced by epithelial-stromal interaction in breast cancer^1^, while Separative cores show elevated expression of metabolic stress-associated genes such as B4GALT2 (related to glycolysis) and DHCR24 (involved in cholesterol biosynthesis)^2^. Furthermore, gene set enrichment analysis revealed that the Mixture group exhibits elevated levels of pathways related to interferon-gamma response, interferon-alpha response, and epithelial-mesenchymal transition (Figure F). In contrast, ROIs displaying a separative pattern demonstrate higher activity in pathways associated with cellular proliferation and metabolism, including MYC targets, E2F targets, G2M checkpoint.

Having observed transcriptomic differences, we aimed to investigate whether cell-cell interactions differ between Separative and Mixture regions. We hypothesized that a receptor gene on sender cells within a tissue region should correlate with the expression of its ligand pair on receiver cells in the same region. To test this, we adapted a receptor-ligand analysis method developed for GeoMx data to predict co-regulated gene pairs, focusing on curated receptor-ligand (RL) pairs. Our analysis revealed that Mixture regions exhibit more receptor-ligand interactions compared to Separative regions (Figure G). Specifically, we identified 47 RL pairs in Mixture regions, whereas only 11 pairs were found in Separative regions. RL pairs specific to Mixture regions were enriched in pathways related to cell adhesion molecules, cytokine-cytokine receptor interactions, and ECM-receptor interactions. Notably, three MHC-T cell marker pairs (HLA-A-CD8A, HLA-F-CD8A, HLA-DRA-CD4) were uniquely highly correlated in Mixture regions.

In conclusion, our results underscore the importance of integrating in situ imaging with spatial transcriptomics for a more accurate and reliable analysis of tumor tissues. Our approach provides critical insights into the tumor microenvironment and cellular interactions, with significant implications for both research and clinical applications in oncology.

## METHODS

### ROICellTrack

The analysis toolkit uses ROI images from the GeoMx DSP platform as the input. In the GeoMx DSP Analysis Suite (version 3.0.0.113 used in this analysis), to output the ROI images, go to the left panel in the slide view, select “Export Image,” choose “ROI Report” in the Export, and then select “TIFF” as the Format. This step will generate square TIFF images with the segmented ROI centered within each image. In our example data, each circular ROI segment is approximately 300 µm in diameter, and the output image is around 600 µm by 600 µm in size. The code provided in the ROICellTrack toolkit then performs three main steps: (1) python code to facilitate automatic image cropping and zooming, removing all content outside of the ROI (including the white margin); (2) python code for cell segmentation based on a deep neural network, which outputs cell counts, locations, and cellular morphological features; and (3) R code for calculation of spatial statistics (e.g., Cross K intersection) based on the cell coordinates and cell labels.

The cell segmentation of the ROI images is based on neural network models implemented in Cellpose v2.2.3. An advantage of Cellpose is its dual functionality, offering both a graphical user interface (GUI) for straightforward parameter optimization and a command-line mode for automated batch processing. We use the pre-trained TissueNet models by setting model_type=‘TN3’. Before running the segmentation, we recommend using the Cellpose GUI to preview the results and optimize the parameters. The two key parameters include ‘cell diameter’ (default of 23 in the pipeline) and ‘flow_threshold’ set to 0.4 by default. For the main segmentation step of cancer cells in our data, we also set channels = [2, 3] to indicate blue nuclei and green-stained cancer cells (PanCK). To help with cell typing after cell segmentation, the visualization of the density of average color intensities (green/red/blue) for each cell and the clustering plots of red and green intensities based on K-Means and Gaussian Mixture Model (GMM) clustering are also performed. In our analysis, we found that GMM better fits the distribution of the red vs. green plot and provides better performance in cell classification. We also discovered that the green density has a very clear bimodal distribution, allowing for easy dichotomization (with a cutoff at 20 in our data). The analysis pipeline will automatically output a snapshot of the original image and cell-annotated segmented images, indicating the total number of cells, the total number of cancer cells, and the proportion of cancer cells. A pandas.DataFrame object will also be saved which will store calculated cell-level metrics, including red intensities, green intensities, blue intensities, number of pixels, perimeters, areas, circularity, and X, Y coordinates.

### Collection of clinical samples

Six formalin-fixed, paraffin-embedded (FFPE) samples from three patients who underwent concomitant radical cystectomy and nephroureterectomy for concurrent upper-tract urothelial carcinoma (UTUC) and urothelial carcinoma of the bladder (UCB) were identified at Moffitt Cancer Center. For each FFPE specimen, a pathologist identified and selected 18 regions of interest (ROIs) based on the lower-tract (LW-Tu) and upper-tract (UP-Tu) locations and the presence of immune cells. Multiplex immunofluorescence (IF) was performed on each FFPE specimen, staining for DNA, Pan-CK, CD20 (B cells), and CD45 (immunocytes). The GeoMx™ Digital Spatial Profiling platform was used to generate RNA profiles from the ROIs on each FFPE slide.

### Spatial clustering

Inspired by Feng et al.’s work ^3^, the cross K AUC score was used to assess the spatial clustering degree between two types of cells – tumors and non-tumors, as defined by ROICellTrack. The Kcross function, implemented in the spatstat R package, was utilized to compare observed patterns with a reference Poisson Process. The AUC score was determined by calculating the area between the curve of the cross K function and the reference pattern.

### Spatial transcriptomics analysis

**Data generation:** Whole transcriptomic profiling and immunostaining were performed using the Nanostring GeoMX®-DSP RNA Assay kit (NanoString Technologies, Seattle, WA) on the BOND RX autostainer (Leica Biosystems, Vista, CA). FFPE tissue sections were baked at 60°C for 60 min before being transferred to the BOND RX (Leica Biosystems). The deparaffinization, antigen retrieval, and hybridization were carried out using the fully automated GeoMX-DSP FFPE RNA Assay Protocol (Nanostring). Subsequently, slides were washed and stained with morphology markers PanCK, CD45, and Syto13 for 2 hrs. ROIs were annotated by a pathologist to include tumor cores, stroma regions, and tumor-stroma margins and were placed on 20X fluorescent images scanned by GeoMx® DSP. The photocleaved oligos from the spatially resolved ROIs in the microplate were quantified using standard next-generation sequencing Illumina® workflows. **Data processing**: Downstream bioinformatics analysis was performed using R packages GeoMxTools and NanoStringNCTools. Sequencing quality was assessed for each segment and probe. A gene-level count matrix was generated by averaging corresponding probes per segment. Segments with very few detected genes (<5% of the total detected genes) would be filtered out, and only genes detected in at least 10% of the ROIs were retained. Following these criteria, data from all 56 ROIs and 11,450 genes were retained. The filtered data were then normalized using upper quartile (Q3) normalization. **Deconvolution analysis**: R package SpatialDecon was used to deconvolve the cellular composition of each ROI. To estimate not only stroma and immune cells but also tumor abundance, we utilized a pre-built reference matrix SafeTME, along with tumor-specific signatures identified from the tumor cores of the current study. The results were visualized by stacked barplot using the R package ComplexHeatmap. **Differential expression test**: The differences in overall transcriptomic landscape across ROIs were examined with UMAP dimension reduction and unsupervised clustering. Differential gene expression between groups of interest was tested with a linear mixed-effect model (LMM). **Receptor-ligand analysis**: A method was developed on GeoMx data to predict receptor-ligand pairs by examining the Pearson correlation between ligand-target gene pairs generated by the R package NicheNetb^4,5^. This method was applied to Mixture and Separative ROIs separately. Specifically, a gene expression matrix was first generated for each type of ROI, and then the pairwise Pearson correlation for any pair of genes was tested. The data were then filtered with a correlation > 0.75, and gene pairs documented in NicheNet that are not based on PPI prediction were kept. Data were visualized using the R package CCPlotR^6^.

---

**Figure 1.**
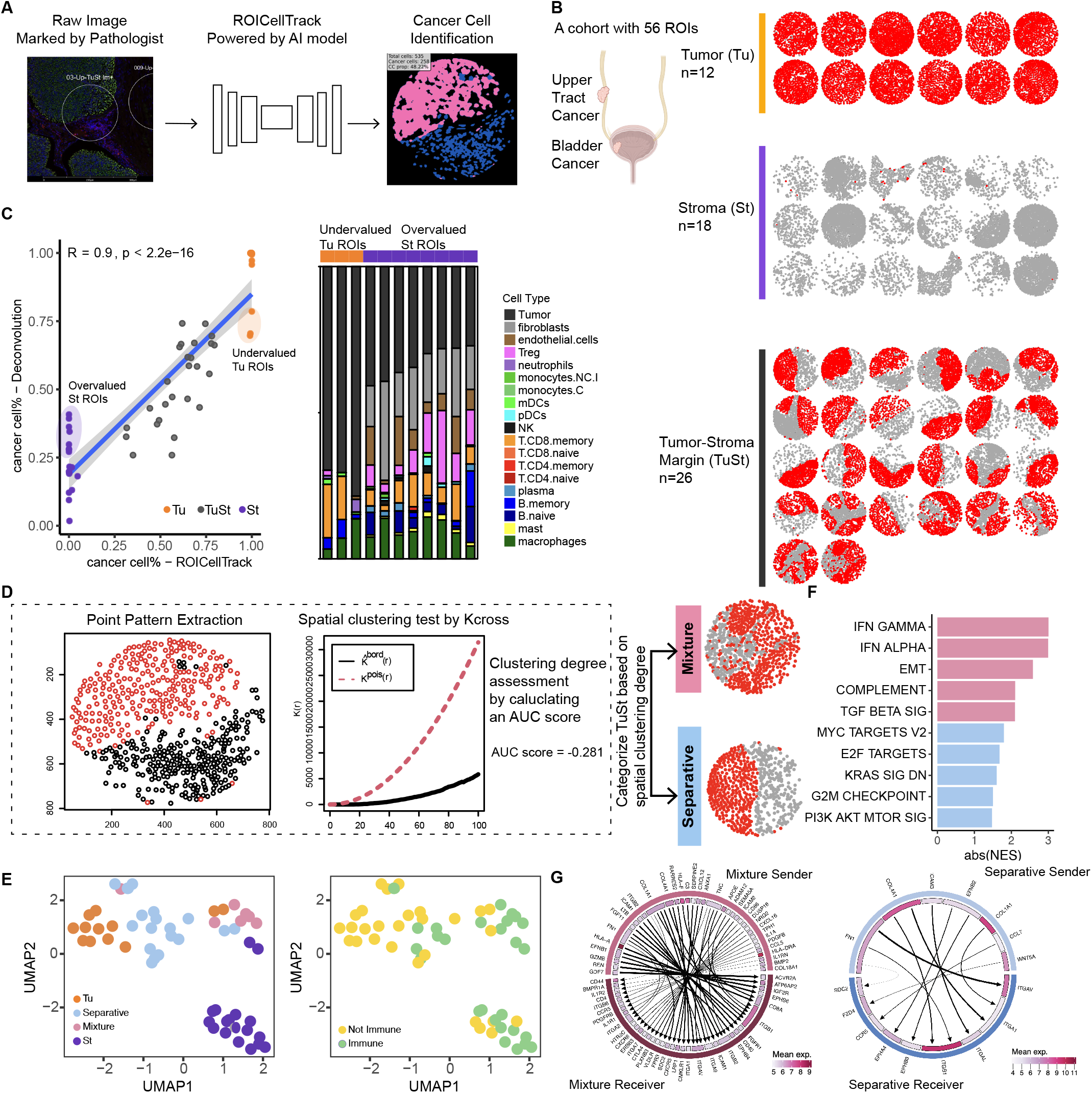
Leveraging spatial transcriptomic profiling with cellular imaging modality. (A) Schematic representation of ROICellTrack, a deep-learning framework designed to identify cancer cells and extract image features. (B) Summary of a bladder cancer cohort with 56 Regions of Interest (ROIs). The ROIs were selected to target three distinct types of regions: Tumor cores (Tu), Stroma (St), and Tumor-Stroma margins (TuSt). Each ROI is visualized using point patterns, with each dot representing an individual cell. Cancer cells are highlighted in red. (C) Comparative performance analysis of ROICellTrack and a gene expression-based cell deconvolution approach. Left panel: Correlation analysis between the two deconvolution methods. Stroma (St) ROIs with overestimated tumor purity and Tumor (Tu) ROIs with underestimated purity are circled. Right panel: Gene expression-based deconvolution results. The predicted cell type composition of each sample is presented as a stacked bar plot. (D) The spatial clustering degree was evaluated for an example ROI. The AUC score was determined by calculating the area between the cross K function curve (black line) and the reference Poisson process (red dashed line). Margin ROIs were grouped based on the clustering degree into Mixture patterns and Separative patterns. One example ROI for each group is provided. (E) UMAP visualization of ROI clustering by region type (left panel) and immune infiltration (right panel). The immune infiltration status is annotated by a pathologist. (F) Molecular pathways enriched in Mixture ROIs (pink) and Separative ROIs (blue). (G) Circos plot showing ligand-receptor pairs within Mixture ROIs (left panel) and Separative ROIs (right panel). Arrows connect gene pairs that are highly correlated, pointing from the ligand to its target gene. The mean normalized expression of each gene is shown in the inner circle.

